# The first reference genome assembly of the Chilean sea fig (*Carpobrotus chilensis*)

**DOI:** 10.64898/2026.06.15.732467

**Authors:** Haein Lee, Carla M. D’Antonio, Soojin V. Yi

**Affiliations:** Department of Ecology Evolution, and Marine Biology, University of California, Santa Barbara, CA 93106, USA; Neuroscience Research Institute, University of California, Santa Barbara, CA 93106, USA; Environmental Studies Program, University of California, Santa Barbara, CA 93106, USA; Department of Molecular, Cellular and Developmental Biology, University of California, Santa Barbara, CA 93106, USA

**Author notes:** Corresponding authors: Carla M. D’Antonio, Soojin V. Yi.

**Keywords:** *Carpobrotus chilensis*, Aizoaceae, genome assembly, PacBio HiFi

## Abstract

*Carpobrotus chilensis* (Chilean sea fig) is a coastal succulent of uncertain origin that has naturalized along the California coast, where it co-occurs and hybridizes with the invasive species, *Carpobrotus edulis.* Despite their ecological importance and widely supported hybridization, genomic resources for this genus remain scarce. Here, we present a draft genome assembly of *C. chilensis* generated from PacBio HiFi long reads. The assembled nuclear genome spans 981.7 Mb across 178 contigs. The contig N50 was 73.0 Mb, and BUSCO completeness was 96.3%. K-mer and SNP-based analyses indicate extremely low heterozygosity (3.4 × 10⁻⁴), reduced genetic diversity in this population. The genome is highly repetitive, with 81.67% of the sequences composed of transposable elements, predominantly long terminal repeat (LTR) retrotransposons. Gene prediction identified 21,744 protein-coding genes, with BUSCO completeness of 95.8%. Comparative analysis with *C. edulis* identified 8,783 single-copy orthologous gene pairs, with a median synonymous substitution rate (dS) of 0.019, indicating low sequence divergence between the two species. This genome assembly provides a foundational resource for investigating the genomic basis of hybridization and invasion in *Carpobrotus*.

## Introduction

*Carpobrotus* species are mat-forming succulents in the family Aizoaceae, most of which are native to South Africa (Hartmann, 2002). While they are traded and planted because they provide societal benefits such as slope stabilization and ornamental use (Campoy et al., 2018), they have also become invasive in many regions of introduction because they spread rapidly beyond planted sites, overgrow native species, alter soil chemistry and change food web relationships (e.g. Molinari et al., 2007; Campoy et al., 2018). Hybridization has been reported among *Carpobrotus* species, both between introduced (Suehs et al., 2004; Campoy et al., 2019; Novoa et al., 2023) and potentially native congeners (Albert et al., 1997; Gallagher et al., 1997), often resulting in extensive hybrid swarms.

Along the California coast, hybridization between a species reported as *C. edulis* (Jepson eFlora, 2026) and *C. chilensis* (Figure 1A–B), a species whose native status remains debated (Bicknell & Mackey, 1988; Novoa et al., 2023), appears to be widespread, resulting in hybrid swarms throughout coastal habitats (Albert et al., 1997; Gallagher et al., 1997). Ongoing hybridization is considered likely to contribute to genetic introgression and the potential loss of genetically distinct *C. chilensis* populations (Albert et al., 1997; Gallagher et al., 1997; Schierenbeck et al., 2005).

**Figure 1.**
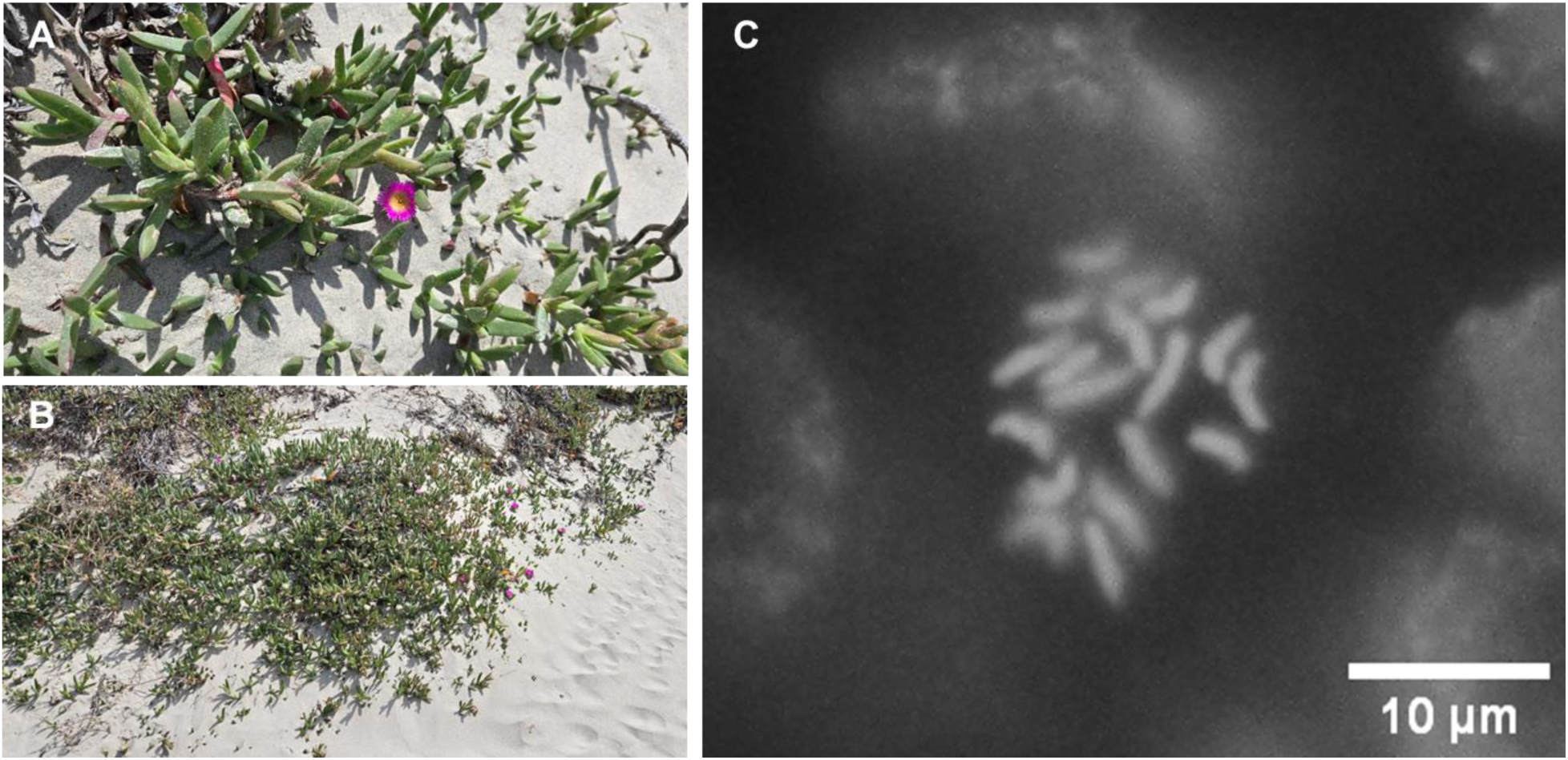
*Carpobrotus chilensis* and its growth habit at San Miguel Island. (A) Representative morphology of *C. chilensis*. (B) Growth habit of *C. chilensis* in its natural coastal habitat on San Miguel Island, Channel Islands National Park, Santa Barbara County, California, USA. (C) Chromosome count of *Carpobrotus chilensis*. Representative chromosome squash showing diploid chromosome number (2n = 18).

Hybridization between *C. edulis* and *C. chilensis* has been investigated using morphological analyses (Albert et al., 1997), allozyme markers (Gallagher et al., 1997), chloroplast DNA (Schierenbeck et al., 2005), and microsatellite-based approaches (Novoa et al., 2023), all of which have provided evidence for introgression in natural populations, but are limited in resolution for characterizing genome-wide patterns of divergence and admixture. As a result, key questions regarding the genomic structure of hybrid populations, the extent and direction of introgression, and the distribution of ancestry across the genome remain unresolved. Addressing these questions requires genome-wide data that enable direct comparison between parental species and their hybrids. Genome resources for the genus *Carpobrotus* remain limited. Recently, a draft reference genome for *C. edulis* has become available (NCBI Assembly GCA_965788445.1). However, the lack of a corresponding genome for *C. chilensis* has restricted genome-wide investigations of divergence and introgression, as well as accurate inference of ancestry in hybrid populations. This asymmetry has constrained comparative analyses between the two species at the whole-genome level.

Here, we present a draft genome assembly of *Carpobrotus chilensis* generated using PacBio HiFi sequencing from an isolated population on San Miguel Island, Channel Islands National Park in Santa Barbara County, California, one of the few locations where *C. chilensis* is abundant and where *C. edulis* has not been reported (Hasenstab-Lehman & Guilliams, 2021), allowing sampling of individuals with minimal expected hybrid influence. This assembly provides a foundational genomic resource for the genus and enables direct comparison between the two parental species, facilitating future investigations of genomic divergence, hybrid ancestry, and introgression dynamics in California coastal populations.

## Materials and methods

### Biological material

Biological material used for genome assembly was collected from a single individual of *C. chilensis* on San Miguel Island (34.032950°N, 120.363932°W) in Channel Islands National Park, Santa Barbara County, California, USA. San Miguel Island was selected because *C. edulis* and hybrid populations are absent, reducing the likelihood of sampling admixed individuals. The collected individual was propagated in a greenhouse, and fresh young leaves were harvested and snap-frozen in liquid nitrogen prior to DNA extraction.

### Chromosome preparation and counting

Actively growing root tips were collected in the morning and pretreated in ice cold water for 24 hours. Samples were fixed in 3:1 (v/v) mixture of ethanol and acetic acid for 24 hours and stored in 70% ethanol at 4°C until use. Root tips were macerated in 1N HCl at 60°C for 10 minutes, stained with 2% acetocarmine for 1 hour, and the apical meristem was excised. Cells were squashed in 1% acetocarmine under a coverslip and used for chromosome observation and counting. Chromosomes were visualized using fluorescence microscopy with a TRITC filter set, taking advantage of the intrinsic fluorescence of carmine-stained chromatin (Stockert et al., 1990).

### High-molecular-weight DNA extraction for PacBio HiFi sequencing

High-molecular-weight (HMW) genomic DNA was extracted from approximately 1 g of young leaves (approximately 1 cm in length) that were stored at −80 °C and used without further sectioning. DNA extraction followed the sorbitol pre-wash and high-salt CTAB protocol described by Inglis et al. (2018), including their recommendations for HMW DNA handling. Several modifications described below were introduced to optimize HMW DNA extraction from succulent *Carpobrotus* tissue. Extraction of HMW DNA from succulent species such as *Carpobrotus* (Aizoaceae) can be particularly challenging due to high levels of polysaccharides and polyphenols that interfere with DNA extraction and downstream applications (Diadema et al., 2003). To reduce the effects of these compounds, sorbitol wash steps were repeated one to three times until the supernatant became sufficiently clear. Frozen tissue was ground in liquid nitrogen using a pre-chilled mortar and pestle until a fine powder was obtained, with intermittent re-freezing during grinding to minimize thawing. All extraction steps, except heat incubation, were performed on ice or at 4 °C. To minimize mechanical shearing, samples were handled gently throughout the protocol, avoiding vortexing and using wide-bore pipette tips for resuspension and transfer. Proteinase K was added during lysis at a final concentration of 0.2 mg/mL, followed by RNase A treatment at a final concentration of 2 mg/mL prior to chloroform:isoamyl alcohol (24:1, v/v) extraction. Chloroform:isoamyl alcohol extraction was performed twice to further improve DNA purity. When visible DNA precipitates were observed, they were carefully transferred using wide-bore pipette tips while retaining a small volume of solution to minimize fragmentation. Following these steps, genomic DNA was resuspended in EB buffer (10 mM Tris-Cl, pH 8.5) and incubated at 4 °C overnight to facilitate dissolution of HMW DNA. When DNA purity did not meet the quality requirements for PacBio library preparation, an additional salt-chloroform purification protocol was used to remove polysaccharides and other contaminants (Flores, 2020). DNA quality and purity were assessed using a spectrophotometer (NanoDrop, Thermo Fisher Scientific, Waltham, MA, USA) by measuring absorbance at 260, 280, and 230 nm, and DNA concentration was quantified using a fluorometer (Qubit, Thermo Fisher Scientific, Waltham, MA, USA). DNA integrity and fragment size distribution were evaluated using 0.8% agarose gel electrophoresis and a Femto Pulse system (Agilent Technologies, Santa Clara, CA, USA), respectively. To minimize DNA fragmentation, HMW DNA was maintained under refrigerated conditions prior to library preparation.

### PacBio HiFi library preparation and sequencing

PacBio HiFi library preparation and sequencing were performed at the UC Davis Genome Center. SMRTbell libraries were constructed from purified genomic DNA using the SMRTbell Prep Kit 3.0 following the manufacturer’s protocol. Gel-based size selection was performed using a LightBench instrument to enrich for long DNA fragments prior to sequencing. Sequencing was carried out on a PacBio Revio platform. Circular consensus sequencing (CCS) reads were generated using the PacBio SMRT Link pipeline to produce highly accurate HiFi reads.

### De novo genome assembly

HiFi reads were assembled de novo using Hifiasm (v0.23.0-r691) with default parameters to generate a primary assembly and alternative haplotigs (Cheng et al., 2021). Redundant haplotypic sequences were further removed using purge_dups (v1.2.6) (Guan et al., 2020). Read-depth statistics were calculated using the purge_dups utilities (pbcstat and calcuts) based on PacBio HiFi read alignments to the assembly, and coverage thresholds for haplotig purging were determined from the read-depth distribution. The resulting purged assembly was used for downstream analyses.

### Contamination screening and organelle separation

To obtain a nuclear genome assembly, putative plastid- and mitochondrial-derived contigs were identified and removed using sequence similarity and read-depth criteria. PacBio HiFi reads were mapped to the purge_dups-filtered primary assembly using minimap2 (Li, 2018), and per-contig sequencing depth was calculated using samtools (Danecek et al., 2021). Reads were also mapped separately to chloroplast and mitochondrial references from the *C. edulis* reference assembly (NCBI Assembly GCA_965788445.1) to estimate organelle-derived read depth. Sequence similarity between assembly contigs and organelle reference genomes was assessed using BLASTN from BLAST+ (Camacho et al., 2009). For each contig, alignment coverage was calculated as the proportion of aligned length relative to the contig length.

Contigs were classified as putative organelle-derived if they satisfied all of the following criteria: (i) at least 50% alignment coverage to plastid or mitochondrial reference genomes, (ii) sequencing depth at least threefold higher than the nuclear median, and (iii) plastid- or mitochondrial-mapped read depth ratios of ≥0.10 or ≥0.05, respectively. To further reduce residual plastid-like contigs, the filtered assembly was additionally screened against the assembled chloroplast genome using BLASTN, and contigs showing ≥99% sequence identity and ≥99% query coverage were removed from the final nuclear assembly.

Following organelle filtering, the assembly was further screened using Tiara (v1.0.3) (Karlicki et al., 2022) to identify residual organelle-derived and other non-nuclear contigs, which were removed from the assembly. The resulting assembly was also screened for foreign contamination using the NCBI Foreign Contamination Screen (FCS-GX, v0.5.5) (Astashyn et al., 2024). Contigs were compared against the NCBI reference database to identify potential non-target sequences based on sequence similarity and taxonomic assignment. No contigs were flagged for removal by FCS-GX.

### Genome characterization

Genome characteristics were estimated from PacBio HiFi reads using k-mer frequency spectra generated with Jellyfish (v2.2.10) at k = 21 (Marçais & Kingsford, 2011), and the resulting k-mer histogram was analyzed using GenomeScope 2.0 with default parameters (Ranallo-Benavidez et al., 2020). Genome-wide heterozygosity was estimated from PacBio HiFi reads mapped to the final nuclear assembly using minimap2. Single nucleotide polymorphisms (SNPs) were identified using bcftools mpileup and call (v1.22) (Danecek et al., 2021), retaining only biallelic sites. Variants were filtered by mapping quality (≥20), base quality (≥20), variant quality (QUAL ≥20), and sequencing depth (5 to 2.5× the mean coverage). Heterozygosity was calculated as the number of heterozygous SNPs divided by the total number of callable bases defined by the same depth thresholds.

### Assembly evaluation

Assembly contiguity statistics, including total assembly size, contig N50, and GC content, were calculated from the contig-level assembly using QUAST (v5.3.0) (Gurevich et al., 2013). Gene space completeness was evaluated using BUSCO (v5.8.3) with the eudicotyledons_odb12 lineage dataset in genome mode (Manni et al., 2021; Tegenfeldt et al., 2025). For k-mer-based assembly evaluation, k-mer databases were generated from PacBio HiFi reads using meryl (k = 21) and used as input for Merqury, from which consensus accuracy (QV) and k-mer completeness were estimated (Rhie et al., 2020). Assembly contiguity and composition were visualized using the snailplot implemented in BlobToolKit (v4.5.1) (Challis et al., 2020).

### Chloroplast genome assembly and annotation

The chloroplast genome of *C. chilensis* was assembled using Oatk (v1.0) (Zhou et al., 2025). The chloroplast genome was annotated using GeSeq with the chloroplast reference set for land plants (Tillich et al., 2017).

### Repeat annotation

Repetitive elements in the *C. chilensis* genome were identified using a de novo and homology-based approach implemented in the Extensive de novo TE Annotator (EDTA, v2.2.2) (Ou et al., 2019). EDTA was run in full annotation mode (--step all) with species parameter set to “others”, allowing detection of a broad range of transposable element (TE) families without lineage-specific priors. To improve TE classification and reduce false positives in coding regions, a non-redundant coding sequence (CDS) set derived from BRAKER3 gene predictions was supplied using the --cds option. TE annotation was enabled using --anno 1, which integrates structural and homology-based evidence to produce a curated TE library and genome-wide TE annotation. To produce a soft-masked genome for downstream analyses, RepeatMasker v4.1.9 was run using the EDTA-derived TE library with the -xsmall option. RepeatMasker was executed with RMBlast (rmblastn v2.14.1+) and default parameters, and repeat annotation statistics were summarized from the resulting .out and .tbl files.

### RNA extraction, library preparation, and sequencing

RNA was isolated from approximately 300 mg of mature leaves that had been snap-frozen in liquid nitrogen and stored at −80 °C using the NucleoSpin RNA Plant and Fungi kit (Macherey-Nagel, Düren, Germany), following the manufacturer’s instructions. RNA concentration and purity were assessed using a spectrophotometer (NanoDrop, Thermo Fisher Scientific, Waltham, MA, USA) by measuring absorbance at 230, 260, and 280 nm. RNA integrity was initially evaluated by electrophoresis on a 1.5% agarose gel and further assessed using a TapeStation system (Agilent Technologies, Santa Clara, CA, USA). RNA-seq libraries were prepared using a strand-specific protocol with poly(A) selection (NEBNext Ultra II Directional RNA Library Prep Kit, New England Biolabs, Ipswich, MA, USA) and sequenced on an Illumina NovaSeq X Plus platform (2 × 150 bp paired-end reads) by Admera Health, yielding approximately 34 million read pairs per sample.

### RNA read processing and alignment

Raw RNA reads were processed using fastp (v1.0.1) to remove adapter sequences and low-quality bases (Chen et al., 2018). Adapter sequences were automatically detected for paired-end reads, and polyG tails were trimmed. Quality trimming was performed using a sliding window approach (window size = 4, mean quality threshold = 20), and reads shorter than 35 bp were discarded. Trimmed reads were aligned to the soft-masked genome assembly using HISAT2 (v2.2.2) with the --dta option (Kim et al., 2019). The resulting alignments were converted to sorted BAM files and indexed using SAMtools (v1.22.1) (Barnett et al., 2011; Li et al., 2009).

### Gene prediction and annotation

For both *C. chilensis* and *C. edulis*, gene prediction was performed using the same annotation pipeline to ensure consistency in downstream comparative analyses. The genome assembly of *C. edulis* was obtained from publicly available resources, and repeat annotation and gene prediction were conducted following the same procedures applied to *C. chilensis*.

Protein-coding genes were predicted using BRAKER (v3.0.8), which integrates GeneMark-ETP and AUGUSTUS in an evidence-guided annotation framework (Brůna et al., 2024; Gabriel et al., 2024; Stanke et al., 2006, 2008). The soft-masked genome generated from EDTA and RepeatMasker was used as input. The coordinate-sorted BAM file derived from RNA-seq alignments was provided as transcriptomic evidence to BRAKER. Protein evidence was generated by combining Viridiplantae OrthoDB v12 protein sequences (Tegenfeldt et al., 2025) with additional protein sequences from *Carpobrotus edulis* (NCBI Assembly GCA_965788445.1). The combined dataset was clustered with CD-HIT at 95% sequence identity to generate a non-redundant protein database (L. Fu et al., 2012), which was used as homology evidence for GeneMark-ETP-guided annotation (Buchfink et al., 2015). BRAKER used GeneMark-ETP to integrate transcriptomic and protein evidence for training, followed by gene prediction with AUGUSTUS. TSEBRA was applied internally to select high-confidence gene models by reconciling predictions from GeneMark-ETP and AUGUSTUS based on extrinsic evidence support (Gabriel et al., 2021).

From the BRAKER annotation, the longest isoform per gene was retained using AGAT (v1.5.1) to generate a non-redundant representative gene set (Dainat et al., 2025). Annotation completeness was evaluated using BUSCO (v5.8.3) in protein mode with the eudicotyledons_odb12 lineage dataset based on this longest isoform set (Manni et al., 2021; Tegenfeldt et al., 2025).

Functional annotation of predicted protein-coding genes was performed using eggNOG-mapper (v2.1.13) (Cantalapiedra et al., 2021; Huerta-Cepas et al., 2019). Protein sequences from the longest isoform set were used as input, and annotations were assigned based on orthology using the eggNOG database. Sequence searches were conducted in DIAMOND mode, and Gene Ontology (GO) terms and KEGG ortholog annotations were retrieved for each gene where available (Buchfink et al., 2021).

### Estimation of substitution rates and identification of putatively selected genes

Orthologs between *C. chilensis* and *C. edulis* were inferred using OrthoFinder (Emms & Kelly, 2019), using amino acid sequence FASTA from the longest predicted isoform of each protein-coding gene. Single-copy orthogroups shared between the two species were extracted, and one-to-one ortholog pairs were retained for downstream analyses. Amino acid sequences for each ortholog pair were aligned using MAFFT (Katoh & Standley, 2013), and the corresponding coding sequences were projected onto the amino acid alignments using PAL2NAL to generate codon alignments (Suyama et al., 2006). Codon alignments were converted to the AXT format required by KaKs_Calculator (Wang et al., 2010), and synonymous (dS) and nonsynonymous (dN) substitution rates were estimated under the YN model (Yang & Nielsen, 2000). Ortholog pairs were filtered to remove unreliable estimates, including pairs with non-finite values, dS values outside the range of 0.001 to 2, dN/dS > 10, amino acid alignments shorter than 50 residues, codon alignments shorter than 150 nucleotides, or internal stop codons. Ortholog pairs with dN/dS > 1 and dS ≥ 0.01 were designated as candidates for putative positive selection. The minimum dS threshold was applied to reduce the influence of inflated dN/dS estimates associated with very low synonymous divergence. Gene Ontology (GO) annotations were obtained from eggNOG-mapper annotations, and GO enrichment analyses were performed using the R package topGO (Alexa & Rahnenführer, 2026). Enrichment was tested separately for biological process, molecular function, and cellular component ontologies using Fisher’s exact test with the weight01 algorithm (Alexa et al., 2006), with all filtered ortholog pairs used as the background set. Terms with p < 0.05 were considered significant.

## Results

### Genome size and characteristics

PacBio HiFi sequencing generated 96.3 Gb of reads from 6.38 million reads, with a mean read length of 15.1 kb and a read N50 of 13.8 kb (Supplementary Figure S1), corresponding to approximately 102× genome coverage for *Carpobrotus chilensis*. A total of 93.7% of bases had quality scores ≥Q30. K-mer spectra showed a single dominant peak at approximately 101× coverage, indicating extremely low heterozygosity (Supplementary Figure S2). SNP-based analysis further estimated a genome-wide heterozygosity of 3.4 × 10⁻⁴. K-mer-based analysis suggested a genome size of approximately 940 Mb (Supplementary Figure S2). Chromosome counting confirmed a diploid complement (2n = 18), supporting that the genome of *C. chilensis* is diploid (Figure 1C).

### Genome assembly and quality

The nuclear genome assembly of *C. chilensis* was 981.7 Mb in length and consisted of 178 contigs, with a contig N50 of 73.0 Mb and an L50 of six contigs (Figure 2 and Table 1). The assembly span was broadly similar to the k-mer-based genome size estimate of approximately 940 Mb (Supplementary Figure S2). The largest contig reached 94.2 Mb. The GC content of the assembly was 41.8%. BUSCO analysis using the eudicotyledons_odb12 dataset recovered 96.3% complete genes (93.8% single-copy and 2.5% duplicated), with 1.8% fragmented and 1.9% missing BUSCOs. Consensus quality value (QV) estimated using Merqury was 65.9, and k-mer completeness was 98.4% (Supplementary Figure S3).

**Figure 2.**
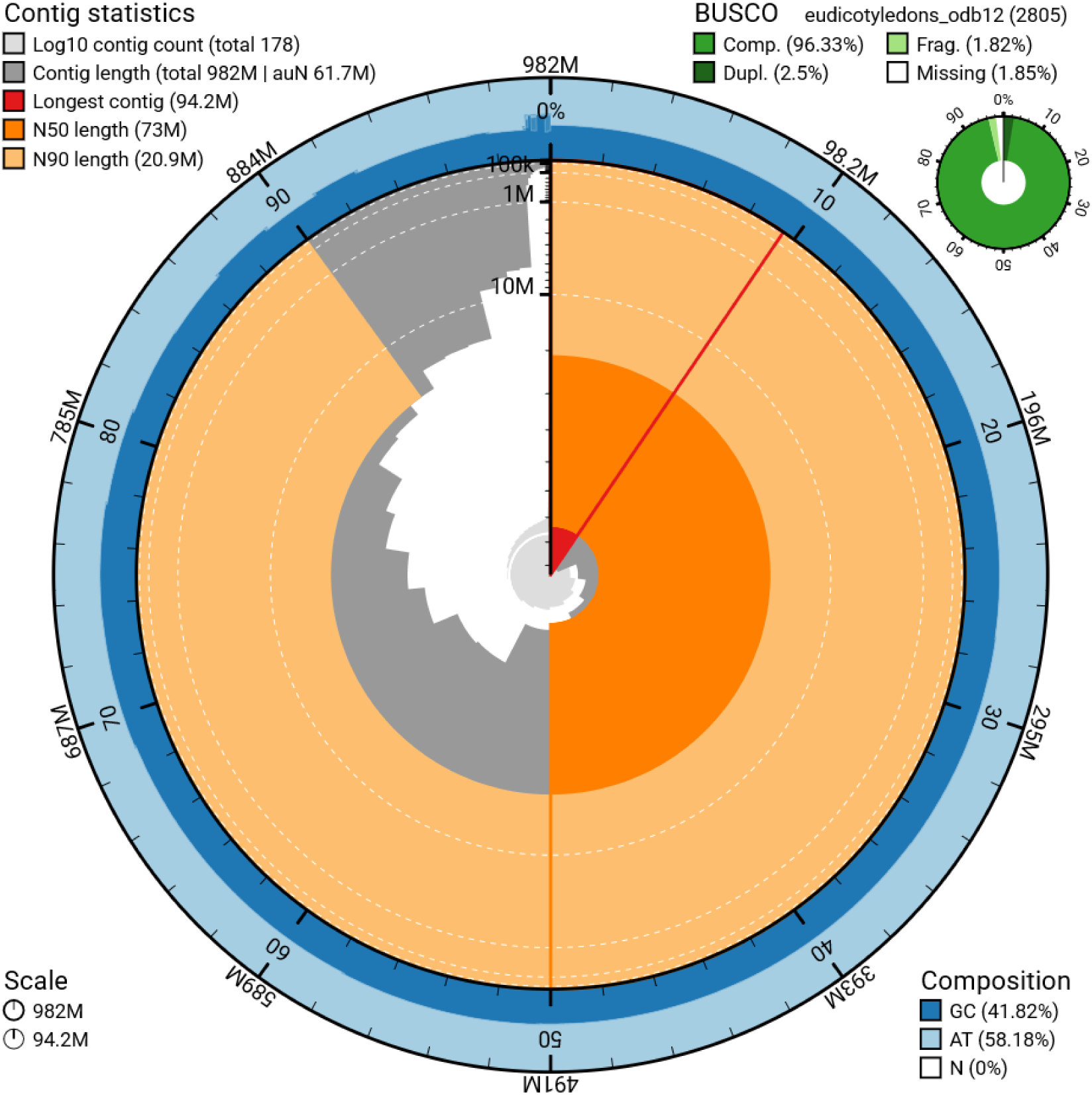
Assembly statistics and quality assessment of the *Carpobrotus chilensis* genome. The snail plot shows contig length distribution, assembly continuity metrics, nucleotide composition, and BUSCO completeness. The assembly spans 981.7 Mb across 178 contigs, with a largest contig of 94.2 Mb and a contig N50 of 73.0 Mb.

**Table 1.**
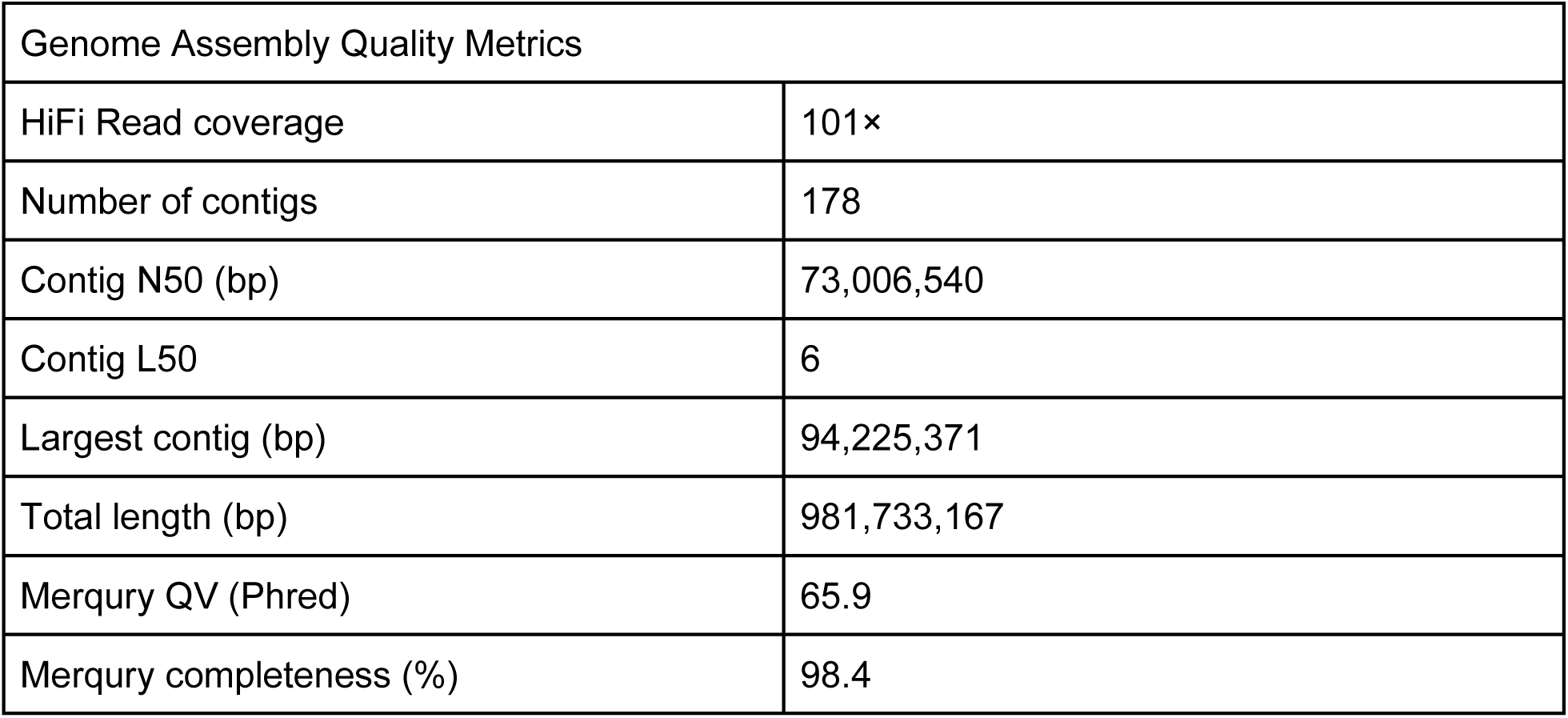
Summary statistics of the *Carpobrotus chilensis* genome assembly.

The chloroplast genome of *C. chilensis* was assembled as a single circular sequence of 148,091 bp, which is similar to the sizes of the chloroplast genomes of *C. edulis* (148,807 bp) and other Aizoaceae plants (NCBI Assembly GCA_965788445.1; W. Fu et al., 2024; Xu, 2019). A total of 105 unique genes were annotated, including 78 protein-coding genes, 23 tRNA genes, and 4 rRNA genes (Supplementary Figure S4).

### Repeat annotation and gene prediction

Repeat annotation suggested a highly repetitive *C. chilensis* genome, with 81.67% of the genome assembly masked as repetitive DNA (Table 2). Interspersed repeats accounted for 80.76% of the genome, with retroelements comprising 70.48%, predominantly long terminal repeat (LTR) elements (69.28%). Among these, *Gypsy* and *Copia* families represented 28.52% and 4.18% of the genome, respectively. DNA transposons accounted for 9.82%, and long interspersed nuclear elements (LINEs) contributed 1.20%. Other repeat categories collectively represented 1.37% of the genome, including unclassified repeats (0.45%), simple repeats (0.81%), and low-complexity regions (0.11%).

**Table 2.** Summary of repetitive elements in the *Carpobrotus chilensis* genome. Classification of transposable elements and other repeat types identified using EDTA and RepeatMasker, including the number of elements, total length occupied, and percentage of the genome.

| Group | Number of elements | Length occupied (bp) | Percentage of sequence (%) |
| --- | --- | --- | --- |
| Retroelements | 942,398 | 691,939,278 | 70.48 |
| LINEs: | 12,910 | 11,797,525 | 1.20 |
| RTE/Bov-B | 256 | 106,754 | 0.01 |
| L1/CIN4 | 12,654 | 11,690,771 | 1.19 |
| LTR elements: | 929,488 | 680,141,753 | 69.28 |
| Ty1/Copia | 72,483 | 41,013,136 | 4.18 |
| Gypsy/DIRS1 | 381,729 | 280,020,217 | 28.52 |
| DNA transposons | 367,195 | 96,448,601 | 9.82 |
| Unclassified: | 29,786 | 4,442,236 | 0.45 |
| Total interspersed repeats: | - | 792,830,115 | 80.76 |
| Simple repeats: | 136,479 | 7,955,335 | 0.81 |
| Low complexity: | 18,882 | 1,035,086 | 0.11 |

Gene prediction identified 21,744 protein-coding genes with 24,921 transcript isoforms (Table 3). BUSCO analysis of the longest isoform protein set recovered 95.8% complete genes (94.0% single-copy and 1.8% duplicated), with 0.8% fragmented and 3.5% missing BUSCOs.

**Table 3.**
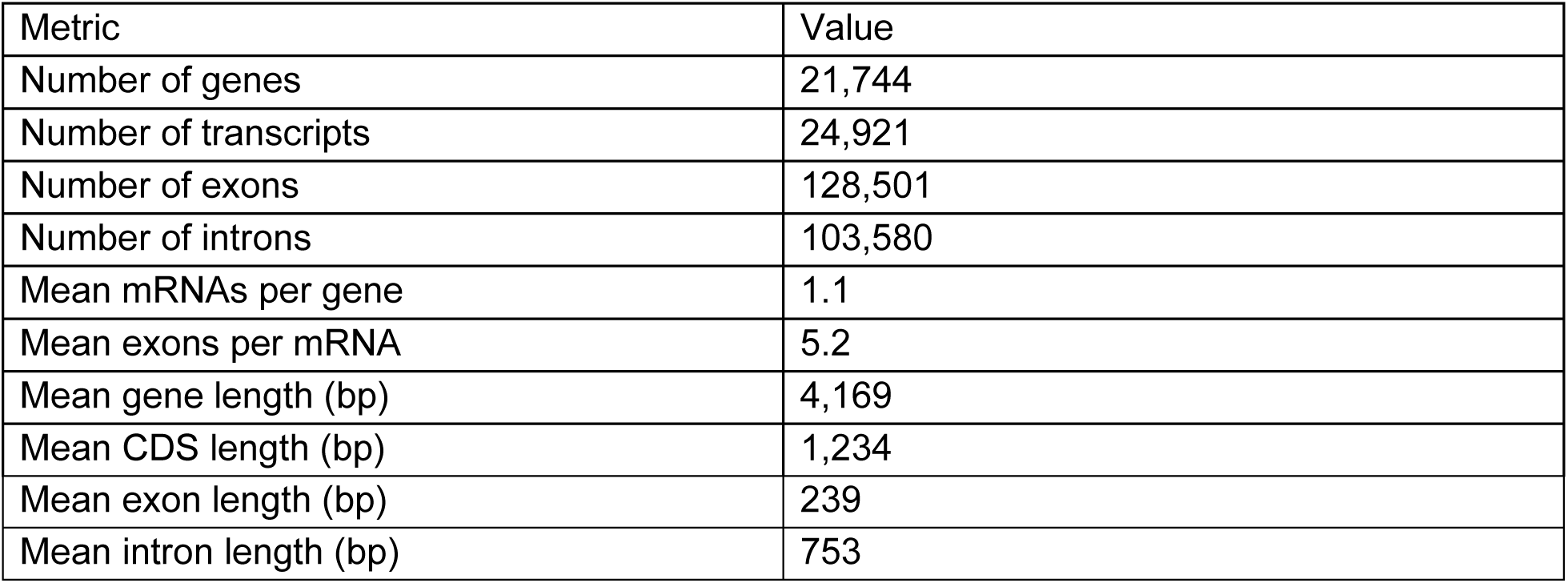
Summary of gene prediction and structural annotation for the *Carpobrotus chilensis* genome. Statistics of predicted protein-coding genes, including gene, transcript, exon, and intron counts, as well as average gene and feature lengths derived from BRAKER3 annotation.

### Divergence of protein-coding genes

A total of 13,254 single-copy one-to-one orthologous gene pairs were identified between *C. chilensis* and *C. edulis*, of which 8,783 were retained after quality filtering. The distribution of synonymous substitution rates (dS) showed a low median value (0.019; mean = 0.038), with the majority of values concentrated below 0.05 (Figure 3). These results indicate low sequence divergence between the two species. The distribution of dN/dS ratios showed a median of 0.290 (mean = 0.392), with most values below 1 and a small number of higher values (Supplementary Figure S5), consistent with purifying selection acting on the majority of orthologous genes.

**Figure 3.**
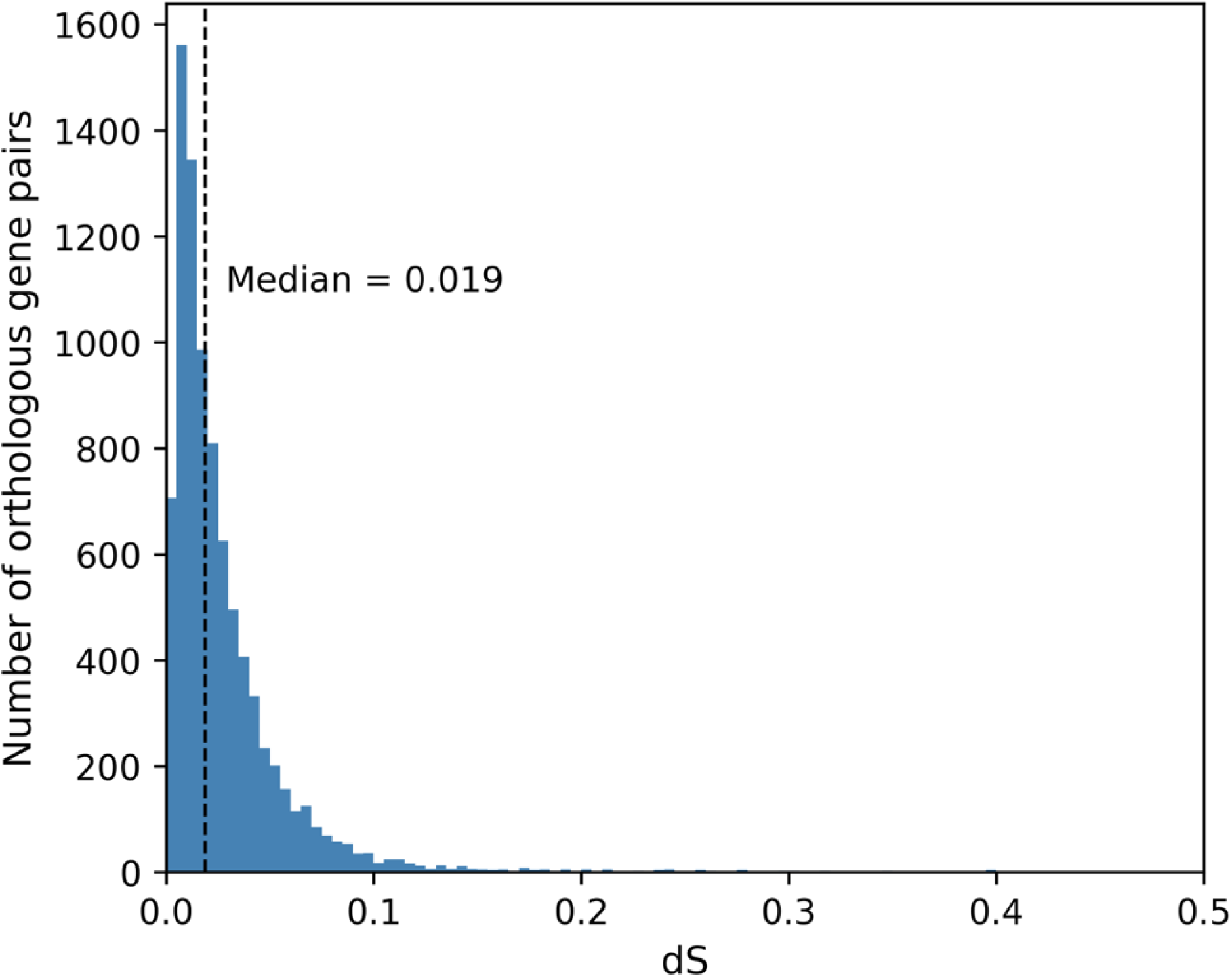
Distribution of synonymous substitution rates (dS) between *Carpobrotus chilensis* and *Carpobrotus edulis*. Frequency distribution of dS values estimated from 8,783 filtered single-copy orthologous gene pairs. The dashed vertical line indicates the median dS value (0.019).

A total of 163 ortholog pairs met the criteria for putative positive selection. Gene Ontology enrichment analysis identified terms associated with defense and signaling processes, including defense response, signal transduction, and cell surface receptor signaling, as well as calcium ion transmembrane transport and leaf morphogenesis (Supplementary Figure S6).

## Discussion

The *C. chilensis* genome assembly presented here represents the first nuclear genome assembly for this species and expands the currently limited genomic resources available for the genus *Carpobrotus*, a globally distributed and ecologically important genus (Campoy et al., 2018; Novoa et al., 2023). Together with the recently published genome of *C. edulis*, this assembly provides genomic references for both putative parental species involved in hybridization along the California coast and establishes a foundation for comparative genomic analyses within the genus.

The assembly also revealed a highly repetitive genome, with 81.67% of the genome composed of transposable elements, predominantly LTR retrotransposons.

A notable characteristic of the *C. chilensis* genome is its extremely low heterozygosity (3.4 × 10⁻⁴). This low level of variation may reflect demographic history, self-fertility (Vilà et al., 1998), clonal propagation, or a combination of these factors. Because the inference of low heterozygosity is based on a single individual, these interpretations remain tentative. Broader sampling across the species’ geographic range will be required to distinguish among demographic and reproductive explanations.

A comparison of protein-coding sequences between *C. edulis* and *C. chilensis* revealed a median dS of 0.019 across 8,783 single-copy ortholog pairs, indicating low synonymous divergence between the two species. This low divergence is consistent with a close evolutionary relationship and may contribute to the widespread hybridization observed in natural populations in California (Albert et al., 1997; Gallagher et al., 1997; Schierenbeck et al., 2005). The predominance of purifying selection across orthologous gene pairs (median dN/dS = 0.290) suggests that protein-coding sequences have remained largely conserved. Nevertheless, the identification of candidate genes under putative positive selection and their associated biological processes illustrates the utility of the *C. chilensis* genome resource for future comparative and evolutionary genomic studies within *Carpobrotus*.

Previous studies of hybridization in *Carpobrotus* have relied on morphological traits, allozyme markers, chloroplast DNA, and microsatellite-based approaches (Albert et al., 1997; Gallagher et al., 1997; Novoa et al., 2023; Schierenbeck et al., 2005), which provide limited resolution for genome-wide analyses of ancestry and introgression. Together with the existing *C. edulis* genome (NCBI Assembly GCA_965788445.1), the *C. chilensis* genome presented here provides a framework for genome-wide analyses of hybridization, such as SNP-based inference of admixture, characterization of introgressed regions, and more precise estimation of divergence between parental species. The chloroplast genome generated in this study provides an additional resource for future phylogeographic and hybridization studies in *Carpobrotus*.

This assembly has several limitations. First, the genome remains at the contig level, which restricts chromosome-scale comparative analyses with the chromosome-level assembly of *C. edulis*. Second, the genome was assembled from a single individual and thus does not capture genetic variation across the species. Third, gene annotation was performed using BRAKER with leaf RNA-seq and protein homology evidence; therefore, genes primarily expressed in other tissues or at other time points may not be fully represented. Despite these limitations, this *C. chilensis* genome provides a valuable resource for future studies of hybridization, divergence, and ecological adaptation in *Carpobrotus*.

## Data Availability

The genome assembly, PacBio HiFi reads, and RNA-seq data generated in this study have been deposited in NCBI under BioProject PRJNA1468704. The genome assembly is available at DDBJ/ENA/GenBank under accession JBZCIG000000000. Genome annotation resources, including structural and functional gene annotations, repeat annotations, and chloroplast genome annotation files, are available through Zenodo (DOI: 10.5281/zenodo.20687645).

The *Carpobrotus edulis* genome assembly used for comparative analyses is publicly available through the International Nucleotide Sequence Database Collaboration (INSDC) under accession GCA_965788445.1 (BioSample: SAMEA110450299), generated by the Darwin Tree of Life project. Public RNA-seq datasets used for gene prediction are available from the European Nucleotide Archive (ENA) under accessions ERR15315144 and ERR15315142.

## Acknowledgements

We thank the Channel Islands National Park for assistance with collection as well as P. Schuyler for transportation to San Miguel Island. We also thank C. Hannah-Bick for assistance growing specimens in the UCSB Greenhouse. We thank the NRI-MCDB Microscopy Facility at UCSB for access to microscopy instrumentation and Ben Lopez for assistance with microscopy. We acknowledge the Darwin Tree of Life Project for generating and making publicly available the genome assembly of *Carpobrotus edulis* used in this study.

