## Supplementary Figures S1-S6 for "The first reference genome assembly of the Chilean sea fig (*Carpobrotus chilensis*)"

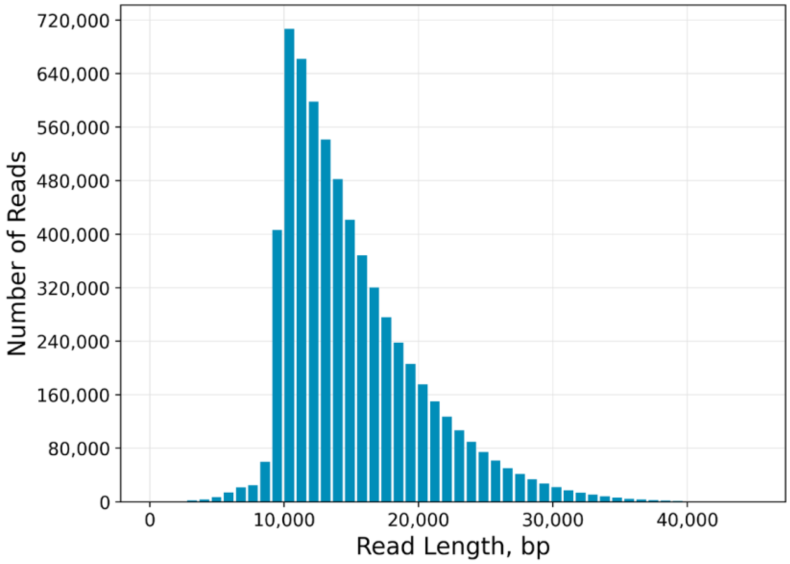


Figure S1. Distribution of PacBio HiFi read lengths for *Carpobrotus chilensis*. Total reads: 6.38 million; read N50: 13.8 kb.

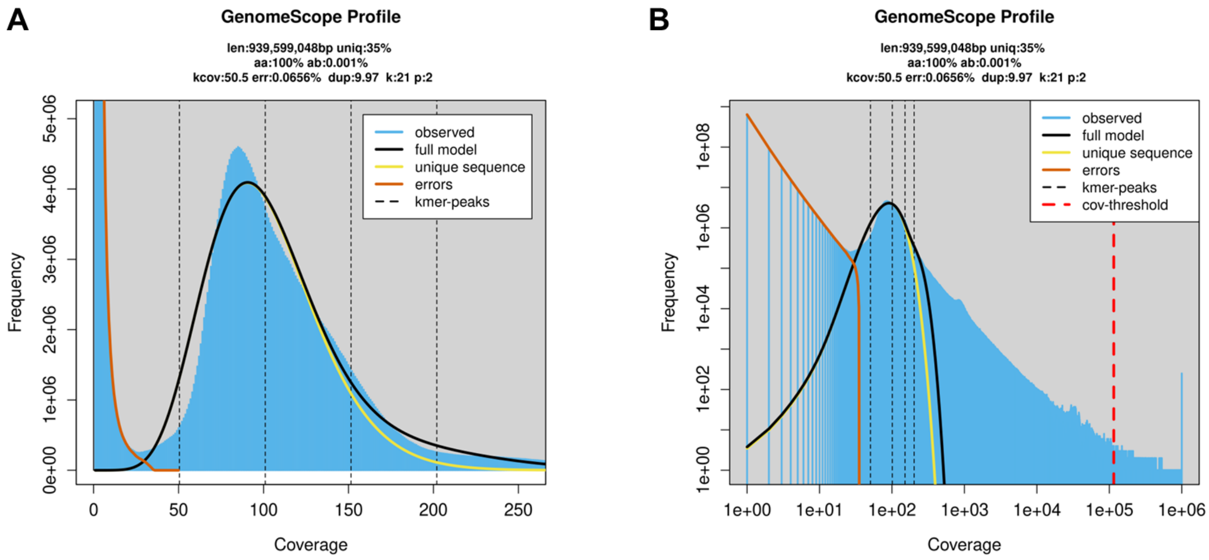


Figure S2. GenomeScope 2.0 analysis of the k-mer frequency distribution of Carpobrotus chilensis. (A) GenomeScope 2.0 profile based on 21-mers shown on a linear scale. (B) The same profile shown on a log scale. Blue bars represent observed k-mer frequencies, and black curves indicate the fitted GenomeScope 2.0 model. Colored curves represent model components: yellow indicates unique sequences, and orange indicates sequencing errors. Vertical dashed lines indicate estimated k-mer coverage peaks; the red dashed line in (B) indicates the coverage threshold.

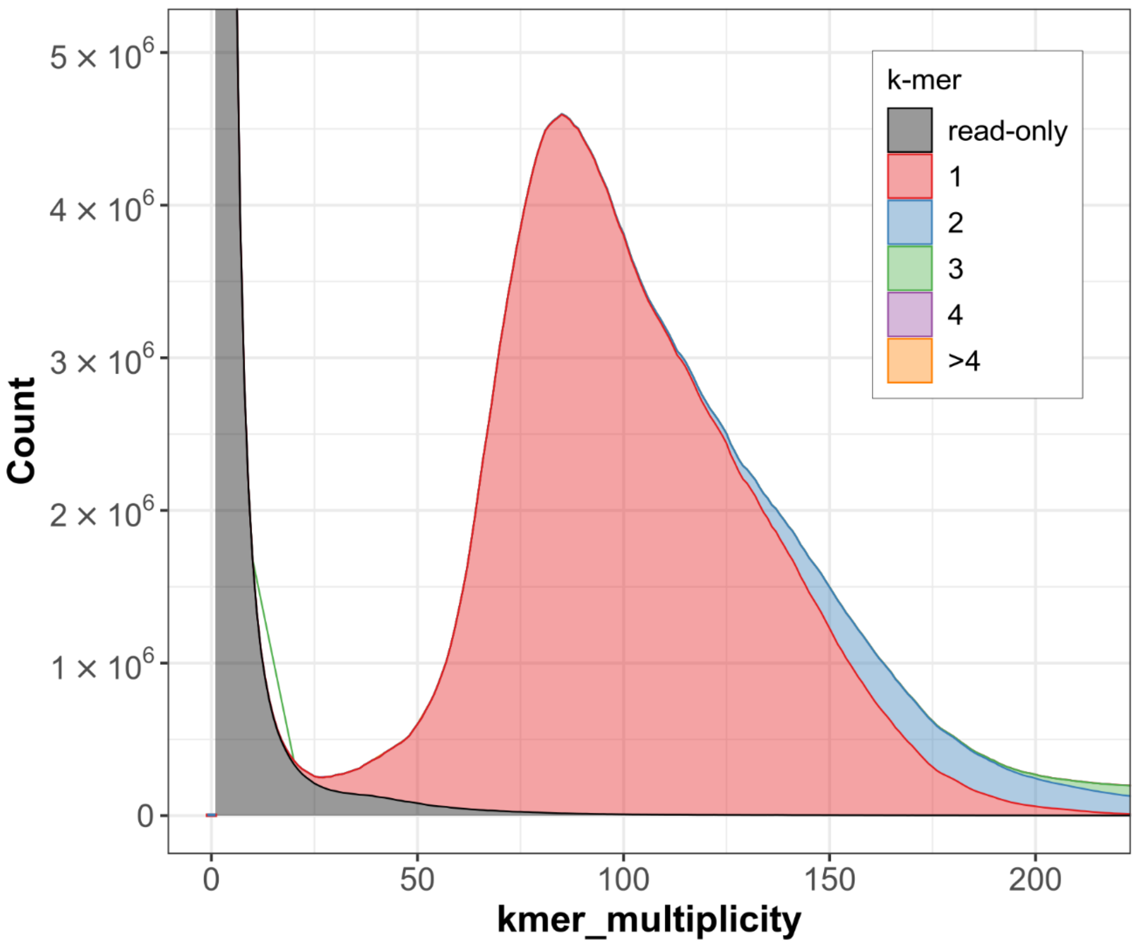


Figure S3. Merqury k-mer multiplicity plot for the *Carpobrotus chilensis* genome assembly. The x-axis represents k-mer multiplicity in the sequencing reads, and the y-axis represents k-mer count. Colors indicate the number of times read-derived k-mers are found in the assembly: gray (read-only, absent from the assembly), red (1 copy), blue (2 copies), green (3 copies), purple (4 copies), and yellow (>4 copies). The dominant red peak at approximately 80–90× multiplicity corresponds to k-mers present as single copies in the assembly. The estimated consensus quality value (QV) is 65.9, and k-mer completeness is 98.4%.

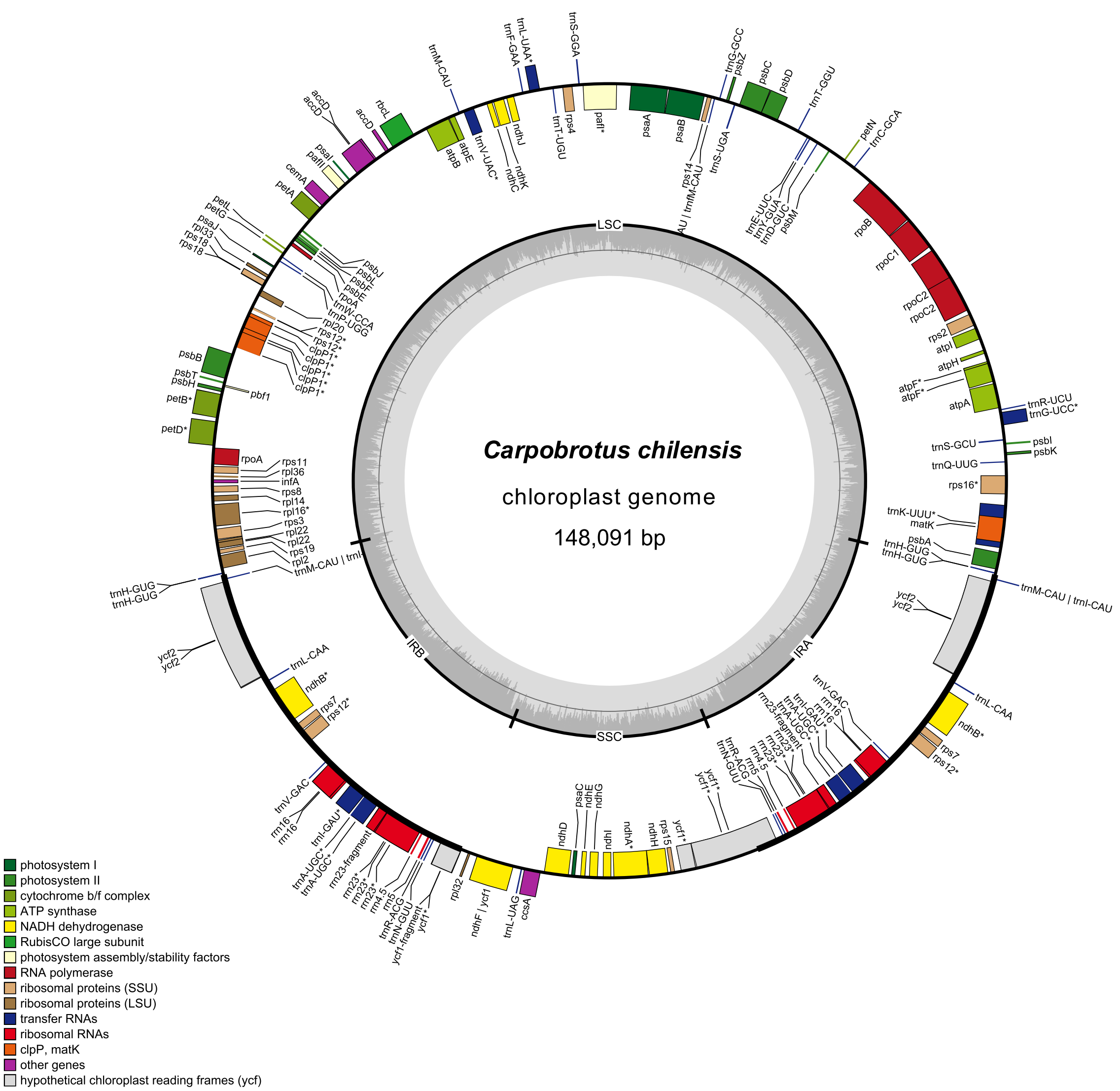


Figure S4. Chloroplast genome map of *Carpobrotus chilensis* (148,091 bp). Genes are color-coded according to their functional categories. The genome structure, including the large single-copy (LSC), small single-copy (SSC), and inverted repeat (IR) regions, is indicated. The gray histogram in the inner circle represents GC content across the genome.

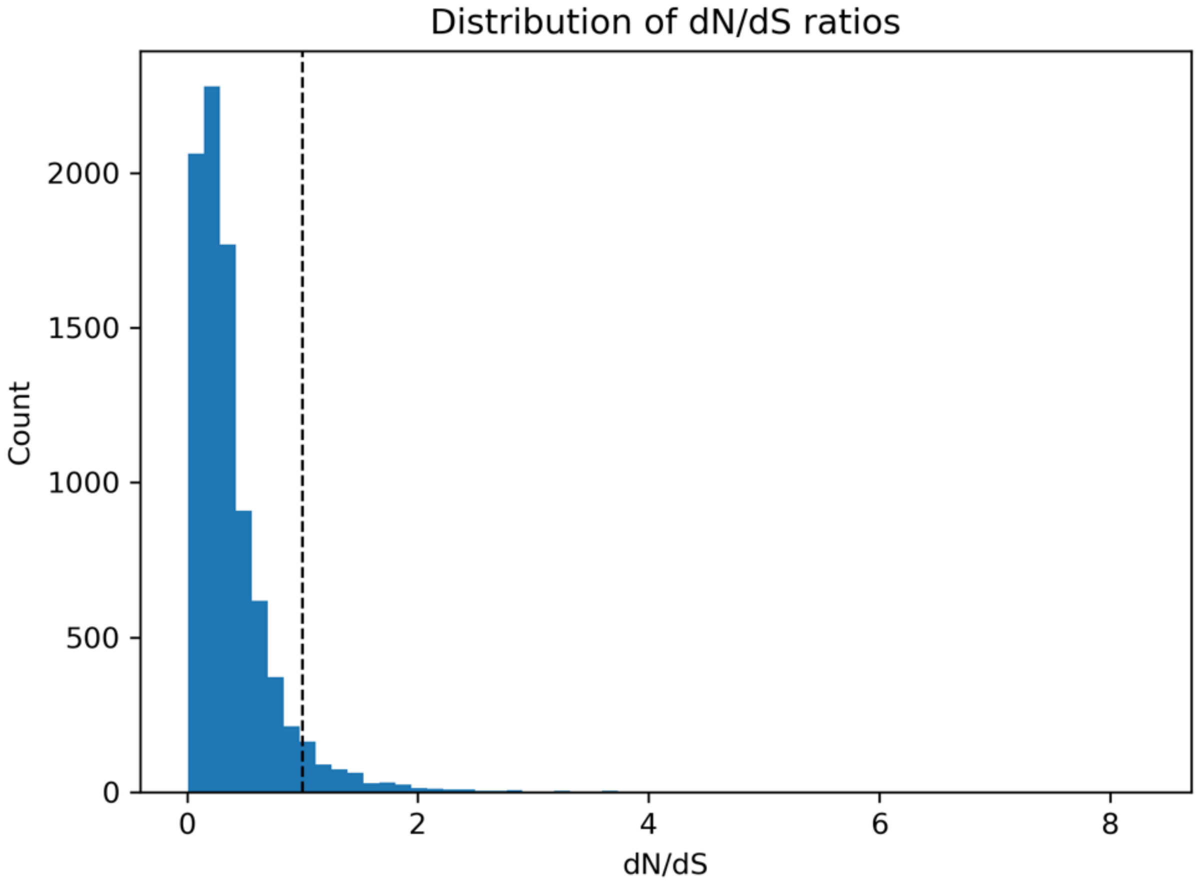


Figure S5. Distribution of dN/dS ratios between *Carpobrotus chilensis* and *Carpobrotus edulis*. Frequency distribution of nonsynonymous to synonymous substitution rate ratios (dN/dS) for filtered single-copy orthologous gene pairs. The dashed vertical line indicates dN/dS = 1.

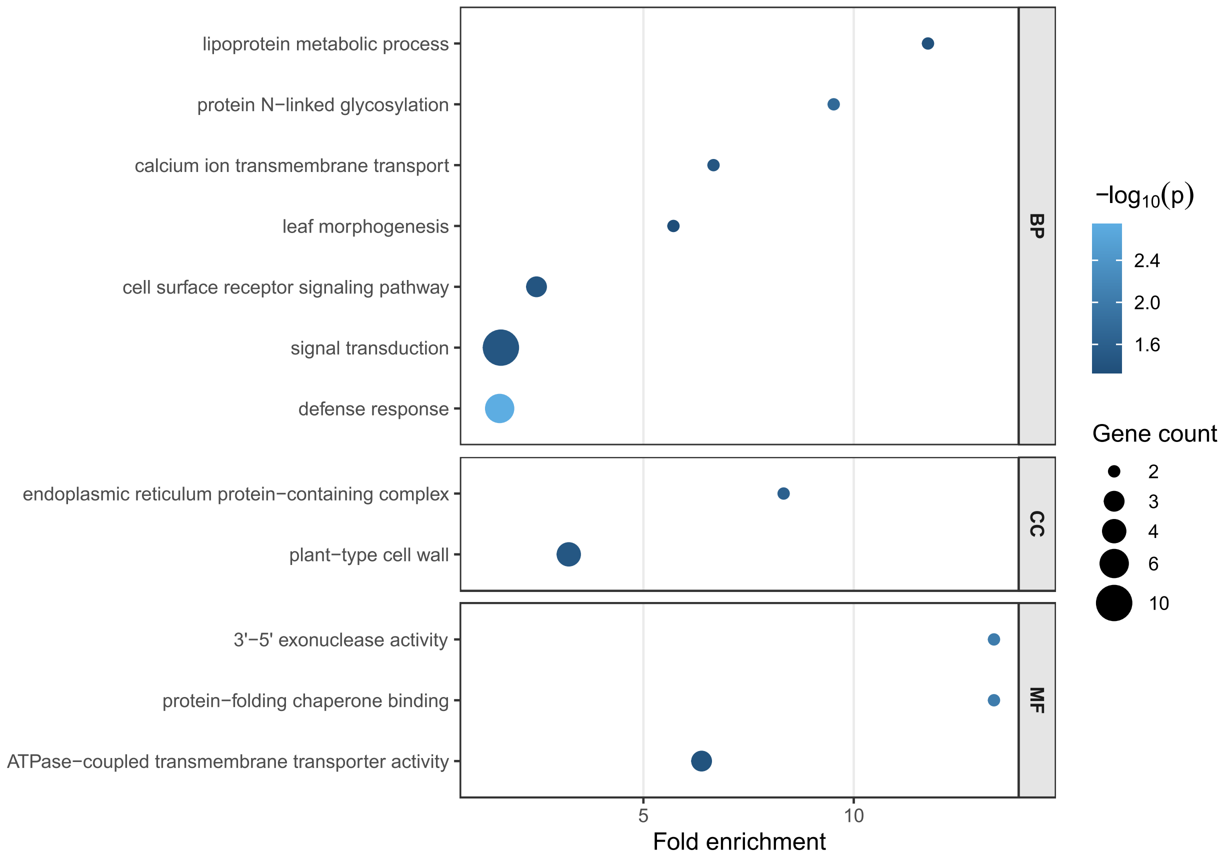


Figure S6. Gene Ontology enrichment analysis of ortholog pairs with dN/dS > 1 and dS ≥ 0.01. Enriched terms from biological process (BP), cellular component (CC), and molecular function (MF) ontologies are shown. The x-axis indicates fold enrichment, bubble size indicates the number of candidate genes associated with each term, and color indicates −log10(P) from the topGO weight01 Fisher test. Only terms with $P < 0.05$ are shown.
